# Cannabichromene is a cannabinoid CB2 receptor agonist

**DOI:** 10.1101/435057

**Authors:** Michael Udoh, Marina Santiago, Steven Devenish, Iain S. McGregor, Mark Connor

**Author notes:** **Correspondence**: Mark Connor, Department of Biomedical Sciences, Macquarie University, Sydney, Australia.

## Abstract

**BACKGROUND:** Cannabichromene (CBC) is one of the most abundant phytocannabinoids in *Cannabis spp*. It has modest anti-nociceptive and anti-inflammatory effects and potentiates some effects of Δ^9^-tetrahydrocannabinol (THC) *in vivo*. How CBC exerts these effects is poorly defined and there is little information about its efficacy at cannabinoid receptors. We sought to determine the functional activity of CBC at CB1 and CB2 receptors.

**EXPERIMENTAL APPROACH:** AtT20 cells stably expressing HA-tagged human CB1 and CB2 receptors were used. Assays of cellular membrane potential and loss of cell surface receptors were performed.

**KEY RESULTS:** CBC activated CB2 but not CB1 receptors to produce a hyperpolarization of AtT20 cells. Activation of CB2 receptors was antagonised by the CB2 antagonist AM630 and sensitive to pertussis toxin. Co-application of CBC reduced activation of CB2 receptors CP55,940, a potent CB1 and CB2 agonist. Continuous CBC application induced loss of cell surface CB2 receptors and desensitisation of the CB2-induced hyperpolarization.

**CONCLUSIONS AND IMPLICATIONS:** Cannabichromene is a selective CB2 receptor agonist displaying higher efficacy than THC in hyperpolarising AtT20 cells. CBC may contribute to the potential therapeutic effectiveness of some cannabis preparations, potentially through CB2-mediated modulation of inflammation.

## Introduction

Cannabichromene (CBC) is one of over 100 phytochemicals collectively referred to as phytocannabinoids that are found in *Cannabis spp* (ElSohly and Gul, 2014). CBC was identified in 1966 and is one of the most abundant phytocannabinoids alongside Δ^9^-tetrahydrocannabinol (THC), cannabidiol (CBD) and cannabinol (CBN) (Izzo et al, 2009; Turner et al, 1980). Evaluation of seized cannabis plants in USA, UK and Australia showed CBC concentrations ranging between 0.05 − 0.3%w/w (Mehmedic et al., 2010; Potter, Clark, & Brown, 2008; Swift et al., 2013). CBC, THC and CBD are directly synthesized from cannabigerolic acid, and share a common 3-pentylphenol ring (Fig 1) (Flores-Sanchez & Verpoorte, 2008). The therapeutic potential of CBC has been evident in several preclinical studies: for example, CBC decreased carrageenan-induced and lipopolysaccharide (LPS)-induced inflammation in rats and mice, respectively (Turner & Elsohly 1981; DeLong et al. 2010); modestly inhibited thermal nociception and potentiated THC anti-nociception in mice (Davis & Hatoum 1983; DeLong et al. 2010). While this may be mediated in part through changes in THC distribution in the mice (DeLong et al. 2010), the pharmacological basis for the *in vivo* actions of CBC remains unclear.

**Figure 1:**
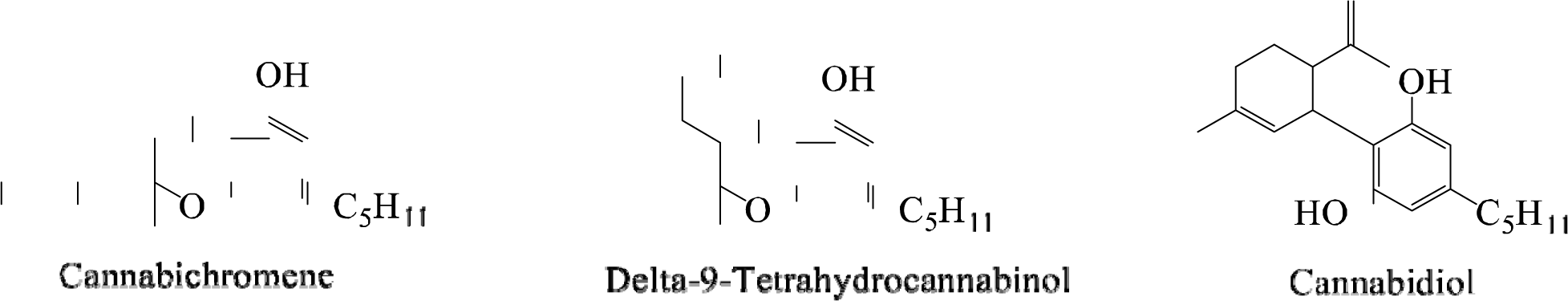
Chemical structures of some phytocannabinoids.

The endocannabinoid system (ECS) comprises the cannabinoid receptors (CB1 and CB2), endogenous agonists including anandamide (AEA) and 2-arachidonoyl-glycerol (2-AG); putative endocannabinoid transporters as well as the enzymes involved in the synthesis and metabolism of endocannabinoids (Iannotti et al, 2016). Cannabinoid receptors are differentially distributed in the body. CB1, the most abundant GPCR in the mammalian brain (Marsicano & Lutz, 1999), is predominantly expressed in the central nervous system, while CB2 is expressed abundantly cells of the immune system and organs such as the spleen. These distributions imply that the activation of these receptors will induce different physiological responses. For example, THC causes a distinctive intoxication via stimulation of the CB1 receptors while stimulation of CB2 receptors does not appear to contribute to the psychoactive effects of cannabis (Deng et al., 2015).

Phytocannabinoids have been found to target individual components of the ECS as well as acting on a range of other receptors and ion channels (Barann et al. 2002; Ryberg et al. 2007). For example, THC activates CB1 and CB2 receptors but also modulates GPR55, 5HT3A receptors and peroxisome proliferator-activated receptor gamma (PPARγ) (Bayewitch et al. 1996; Pertwee 1999; Barann et al. 2002; O’Sullivan et al. 2005; Lauckner et al. 2008). CBD is reported to increase anandamide levels by inhibiting the enzyme FAAH; act as a negative allosteric modulator of CB1 receptors; antagonise GPR55 receptors, activate TRPV2 receptors and like THC, modulate T-type calcium channels(Ross et al., 2008; De Petrocellis et al., 2011; Laprairie et al., 2013). The less prevalent phytocannabinoids tetrahydrocannabivarin (THCV), has been reported to be a low efficacy CB2 agonist, and high potency TRPV1 and TRPA1 agonist (De Petrocellis et al., 2011) while cannabinol (CBN) appears to be an agonist of CB1, CB2 receptors and TRPA1 channels (Bolognini et al., 2010; De Petrocellis et al., 2011; Rhee et al., 1997).

CBC has been reported to be a low affinity CB1/CB2 ligand in binding assays conducted on human receptors expressed in insect cells, and it also activates rat TRPA1 channels (De Petrocellis et al., 2011; Rosenthaler et al., 2014). However, receptor binding does not provide information about ligand efficacy, and whether CBC has efficacy at either receptor remains unresolved. In this study, we sought to characterise the action of CBC at human CB1 and CB2 receptors. To do this, we used an in vitro assay of K channel activation in intact AtT-20 cells that we have used extensively to characterize the activity of cannabinoids at CB1 and CB2 receptors (Banister et al. 2016; Redmond et al. 2016; Soethoudt et al. 2017; Longworth et al. 2017). We find that CBC is an agonist at CB2 but not CB1 in this assay.

## Materials and Methods

### Compounds

CBC was synthesized according to the method of Lee & Wang (2005). Olivetol (1.80 g, 10 mmol) and citral (1.83 g, 12 mmol) were dissolved with stirring in toluene (100 mL), followed by the addition of ethylenediamine (267 uL, 240 mg, 4 mmol) and acetic acid (458 uL, 480 mg, 8 mmol). The mixture was refluxed for 6 h and concentrated under vacuum. The residue was dissolved in dichloromethane (DCM), washed with water and brine, filtered through a plug of silica, and concentrated. Column chromatography was performed multiple times, as separate runs utilising hexane with DCM (gradient from 5:1 to 1:1) and hexane with ethyl acetate or acetone (preferably acetone; gradient from 67:1 to 50:1) were necessary to remove impurities. Cannabichromene was afforded as a pale orange oil (1.20 g, 3.8 mmol, 38%), which darkened upon exposure to air and light. 1HNMR (CDCl3, 400 MHz) δ 6.66 (1H, d, J = 9.9 Hz), 6.29 (1H, s), 6.14 (1H, s), 5.51 (1H, d, J = 9.9 Hz), 5.37 (1H, s), 5.12 (1H, t, J = 6.9 Hz), 2.44 (2H, t, J = 7.67), 2.22-2.07 (2H, m), 1.81-1.66 (2H, m), 1.69 (3H, s), 1.60 (3H, s), 1.59-1.52 (2H, m), 1.41 (3H, s), 1.37-1.25 (4H, m), 0.90 (3H, t, J = 6.6 Hz); 13CNMR (CDCl3, 100 MHz) δ 153.9, 151.1, 144.8, 131.6, 127.2, 124.2, 116.9, 109.1, 107.9, 107.1, 78.3, 41.0, 35.9, 31.5, 30.6, 26.2, 25.7, 22.7, 22.5, 17.6, 14.0.

THC was obtained from THCPharm (Frankfurt, Germany) while CP55,940, SR141716 and AM630 were purchased from Cayman Chemical (Michigan, USA). All drugs were prepared as stock solutions in DMSO and diluted using a 0.01% bovine serum albumin (BSA, Sigma, Castle Hill, Australia) in HEPES-buffered low potassium Hanks Balanced Salt Solution (HBSS). HBSS comprises (mM) NaCl 145, HEPES 22, Na_2_HPO_4_ 0.338, NaHCO_3_ 4.17, KH_2_PO_4_ 0.441, MgSO_4_ 0.407, MgCl_2_ 0.493, Glucose 5.56, CaCl_2_ 1.26; (pH7.4, Osmolarity 315±15mosmol). Final DMSO concentration was 0.1%.

### Cell Culture

Mouse pituitary tumour AtT20 FlpIn cells stably transfected with HA-tagged human CB1 (AtT20-CB1) and human CB2 (AtT20-CB2) receptors (Alexander et al. 2017; Banister, et al. 2016) were used. Tissue culture media and reagents were from Thermo Fisher Scientific, (Massachusetts, USA) or Sigma-Aldrich (Castle Hill, Australia). Tissue culture wares were sourced from Corning (Corning, NY, USA) or Becton Dickinson (North Ryde, Australia). Cells were cultured in T75 flasks using Dulbecco’s modified Eagle’s medium (DMEM) supplemented with 10% foetal bovine serum (FBS) and 1% penicillin-streptomycin (100U/ml) and incubated in a humidified atmosphere with 5% CO_2_ at 37°C. Zeocin (100 µg/ml, Invivogen, California, USA), and hygromycin (80u µg/ml, Invivogen) were used to select wild-type and transfected AtT20 cells, respectively. Cells were passaged at 80% confluency and used for assays at above 90% confluency, for up to 15 passages. For experiments, AtT20 cells were resuspended in Leibovitz’s L-15 media containing 1% FBS, 1% P/S and 15mM glucose. 90 µL of the resuspended cells were plated in a black-walled, 96-well microplate (Corning, NY, USA) and incubated overnight in humidified air at 37°C. For experiments involving pertussis toxin treatment (PTX, Hello Bio, Bristol, UK), 200 ng/ml PTX was added to the L-15 cell suspension.

### Membrane Potential Assay

In this assay, a reduction in fluorescence is indicative of cellular hyperpolarisation. Changes in the membrane potential of cells were measured using a FLIPR Membrane Potential Assay kit (blue), and a FlexStation 3 Microplate Reader (both from Molecular Devices, Sunnyvale, CA). The dye was diluted to 50% of the manufacturers recommended concentration using HBSS. Dye (90 µL) was loaded into each well of the plate and incubated for 1 h at 37°C prior to testing. The FlexStation 3 recorded fluorescence at 2 sec intervals (□_excitation_= 530, □_emission_= 565), and drugs were added after an initial 2 min of baseline reading. The volume of each drug addition was 20 µL, and when two drug additions were made, each drug concentration was adjusted to accommodate the change in final volume. The cellular response to the drug is presented as a percentage change in fluorescence from baseline after subtraction of the change produced by vehicle addition. The change in fluorescence was then normalized to the change in fluorescence due to 1 µM CP55,940 (a high efficacy, non-selective CB1 and CB2 receptor agonist) (Banister et al. 2016) to allow more ready comparison across experiments. Concentration-response curves were fitted to a 4-parameter sigmoidal dose-response curve in Graph Pad Prism (Version 6 GraphPad Software Inc, CA, USA) to derive EC_50_ and Emax.

### Receptor Internalisation Assay

Changes in cell surface CB2 receptors were determined by whole cell enzyme-immunosorbent assay (ELISA). Cells in L-15 media were seeded at 80,000 cells per well in a Poly-D-lysine (Sigma, Castle Hill, Australia) coated, black walled, clear bottom 96-well plate, and incubated for 18 h at 37°C in humidified air. After incubation, cells were treated with the drug of interest. Reported drug concentrations are final concentrations. For one drug treatment, the volume of cells in L-15 and compounds were mixed in a 1:1 ratio. For two drug treatments, the volume of cells in L-15, drug A and drug B were added in ratio 9:9:2. Following drug treatment, receptor trafficking was inhibited by placing cells on ice. Cells were then fixed with 4% paraformaldehyde (PFA) for 15 min. Fixed cells were washed three times with 100 µL Phosphate Buffered Solution (PBS) and blocked with 1% BSA in PBS for 1 h at room temperature. Alexa Fluor^®^ 488 anti-HA Epitope Tag Antibody (Biolegend, UK), diluted to 1:250 with blocking solution, was incubated with the cells at 4°C for 18 h. Cells were then washed three times with 100 µL PBS followed by the addition of 50 µL PBS for the quantification of fluorescence intensity using PHERAstar plate reader (BMG Labtech, Germany). Loss of cell surface receptor was calculated as the percentage decrease in fluorescence intensity after the subtraction of background fluorescence (the fluorescence of wild-type AtT20 cells incubated with the Anti-HA antibody, as above). The background fluorescence was 50 ± 3% of total fluorescence in CB2 expressing cells.

### Data Analysis

All statistical analyses were conducted with GraphPad Prism, in line with the recommendations on experimental design and analysis in pharmacology (Curtis et al., 2015). Data are reported as Mean ± SEM. Two-tailed, unpaired *t*-tests were used to compare two data points and one-way ANOVA for more than two data points. *P*-values < 0.05 were considered statistically significant and indicated with * graphically. Unless otherwise stated, five independent replicates were performed for each experiment.

### Nomenclature of Targets and Ligands

Key protein targets and ligands in this article are hyperlinked to corresponding entries in http://www.guidetopharmacology.org the common portal of data from the IUPHAR/BPS Guide to PHARMACOLOGY 2017/2018 (Alexander et al., 2017)

## Results

### CBC evokes cellular hyperpolarisation via CB2 but not CB1 receptors

CP55,940, a non-selective CB1/CB2 receptor agonist, produced a concentration-dependent decrease in fluorescence in CB1 cells, with a pEC_50_ of 7.8 ± 0.1 (Fig. 2A). THC also evoked membrane hyperpolarisation but with a lower efficacy and potency (pEC50 of 6.6 ± 0.2; Emax of 53 ± 3% of CP55,940) (Fig 2A). CBC did not hyperpolarize CB1 cells, with a negligible change in fluorescence of 2 ± 1% at 30 µM (Fig. 2A-B).

**Figure 2.**
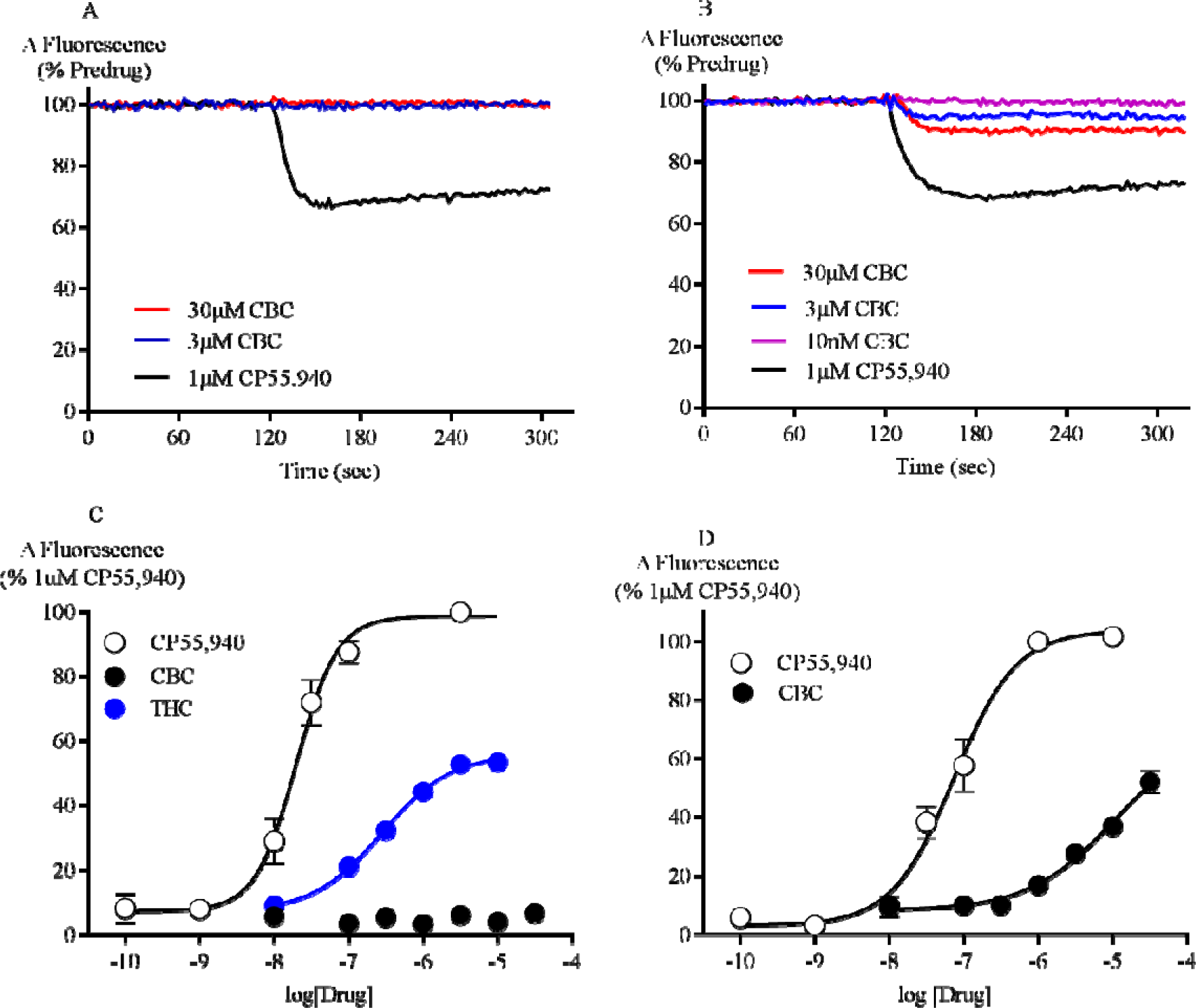
CBC activates CB2 but not CB1 receptors. (A) Representative traces of changes in fluorescence, due to CBC and 1 µM CP55,940-induced hyperpolarisation in (A) AtT20 CB1 cells (B) AtT20 CB2 cells. Drugs were added after 120sec of baseline reading and read over 300sec. (C) Concentration-response curves of CP55,940, THC and CBC in AtT20 CB1 cells. (D) Concentration-response curves of CP55,940 and CBC in AtT20 CB2 cells. Results are expressed as mean ± SEM (n=5-8), after normalization to 1 µM CP55,940 hyperpolarisation.

In CB2 cells, CP55,940 produced a concentration-dependent decrease in fluorescence (pEC50 of 7.1 ± 0.1) (Fig. 2C). The maximum effect of THC (10 µM) was 27 ± 6% of CP55,940. In contrast to AtT20-CB1 cells, application of CBC to AtT20-CB2 cells resulted in a significant hyperpolarisation, reaching a maximum of 52 ± 4% of the maximum effect of CP55,940 at the highest concentration of CBC tested (30 µM, Fig. 2C-D). 30 µM CBC produced a negligible change in fluorescence when applied to wild-type AtT20 cells (2 ± 0.4%).

### CBC-induced hyperpolarisation is blocked by AM630 and is pertussis sensitive

Pre-treatment of AtT20-CB2 cells with AM630 (3 µM, 5 min), a CB2 receptor selective antagonist (Ignatowska-Jankowska, Jankowski, & Swiergiel, 2011), inhibited the 10 µM CBC response by 93 ± 6% compared to vehicle pre-treated cells (Fig. 3A-B). AM630 produced a similar inhibition of responses to CP55,940 (300 nM, Fig. 3B). Overnight incubation of CB2 cells with 200 ng/ml pertussis toxin also abolished responses to 10 µM CBC (8 ± 4% of CP 55,940) and 1 µM CP55,940 (11 ± 4%)

**Figure 3.**
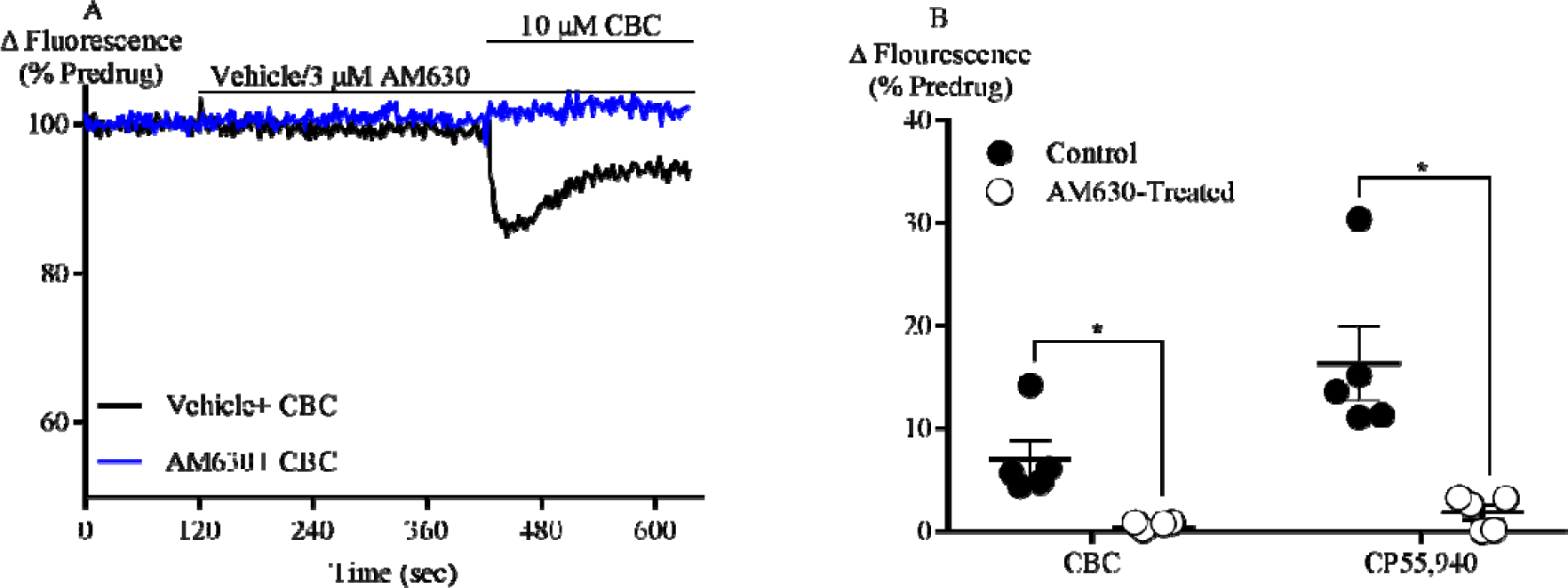
CBC activation of CB2 receptors is blocked by AM630. (A) A representative trace of change in fluorescence of AtT20-CB2 cells after 5 mins pre-treatment with vehicle and 3 µM AM630, followed by the addition of 10 µM CBC. (B) Responses to CBC (10 µM) and CP55,940 (300 nM) in AtT20-CB2 cells with or without pre-incubation of AM630 (3 µM) for 5 mins. Results are expressed as mean ± SEM (*n*=5).

### CBC antagonises CP55,940 in CB2 receptors, but not CB1 receptors

The effect of CBC on responses to CP55,940 and THC was investigated by pre-incubating cells with CBC (10 µM, 5 mins). In CB1 cells, CBC did not significantly affect the subsequent response to either CP55,940 (100 nM) or THC (10 µM) (Fig. 4A-B). In contrast, the CP55,940 (300 nM) response in CB2 cells was significantly inhibited by 44 ± 5% (Fig. 4C & 4E).

**Figure 4.**
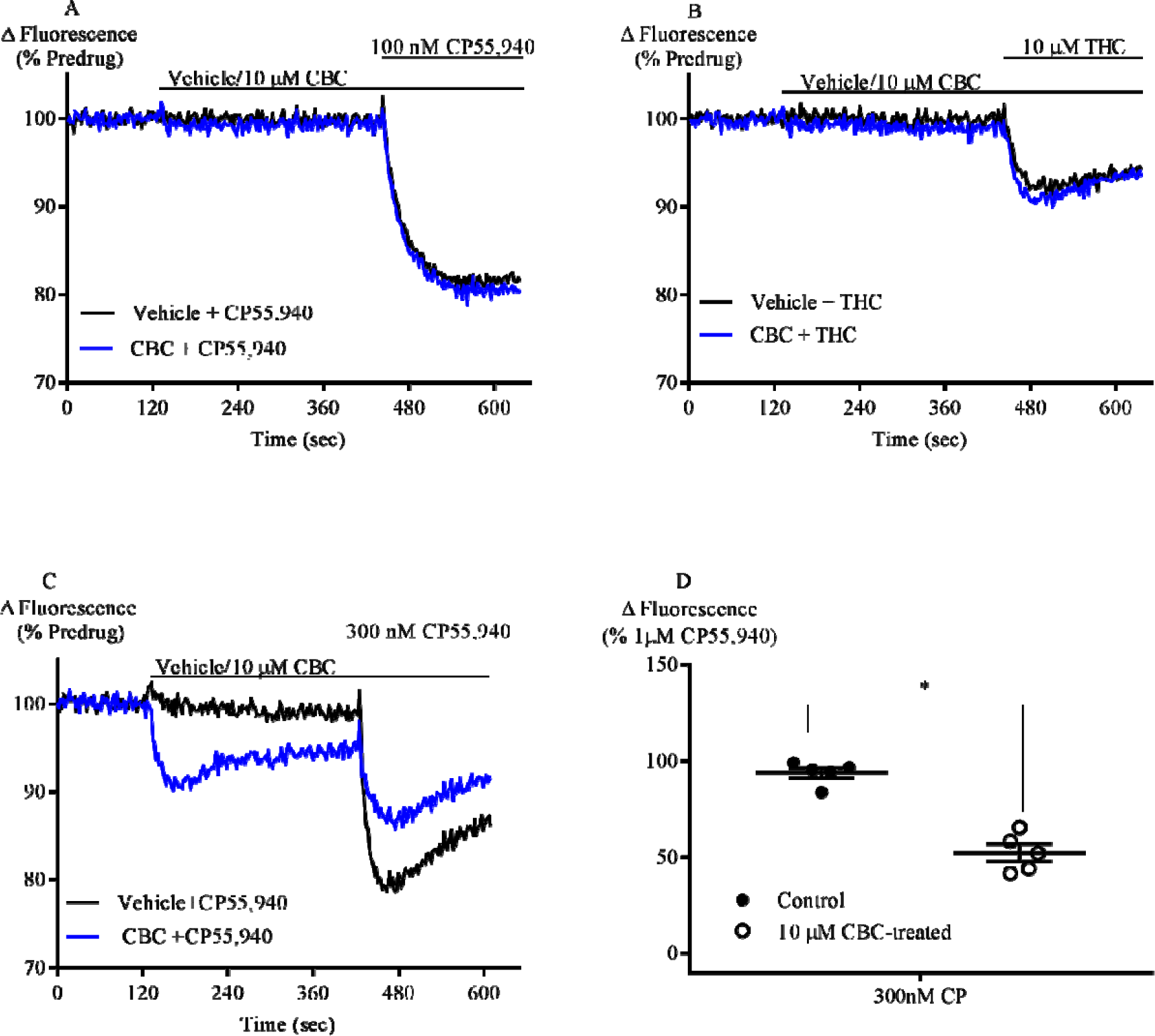
CBC antagonises CP55,940 and THC response in CB2 cells. Representative traces of the effect of CBC (10 µM) on (A) CP55,940 (100 nM) on fluorescence in AtT20-CB1 cells loaded with a membrane potential-sensitive dye. (B) THC (10 µM) hyperpolarisation in AtT20-CB1. (C) CP55,940 (300 nM) hyperpolarisation in AtT20-CB2 cells. After 2mins baseline reading, cells were pre-treated with vehicle or 10 µM CBC for 5 mins, followed by the addition CP55,940 (D) Summary data of the effect of 10 µM CBC on 300 nM CP55,940 in AtT20-CB2 cells. Results are expressed as mean ± SEM after normalization to 1 µM CP55,940 hyperpolarisation (*n*=5) *(Note: Truncated axes).*

### CBC treatment causes cell surface loss of CB2 receptors

CB2 receptors undergo agonist-induced loss of surface receptors following prolonged stimulation (Grimsey et al, 2011). We found that 1 µM CP55,940 internalized CB2 surface receptors to 59 ± 3% of basal surface level (BSL) after 30 min treatment. CBC also internalized CB2 surface receptors (10 µM, 77 ± 5%; 30 µM, 71 ± 3%). When CB2 cells were pre-treated with AM630 (3 µM, 5 mins), 10 µM CBC did not produce significant loss of surface CB2 receptors after 30 mins treatment (108 ± 7% of BSL) (Fig. 5A). The amount of cell surface receptors did not change when AM630 (3 µM, 5 mins) only was used to treat the cells (97 ± 8% of BSL). To assess the possible role of G protein receptor kinases in CBC-mediated receptor internalisation, we pre-treated cells with Compound 101 (10 µM), a GRK2/3 kinase inhibitor (Lowe et al., 2015), for 1 h and then challenged them with CBC. There was no significant change in 10 µM CBC internalisation of CB2 surface receptors following Compound 101 pre-treatment (Fig. 5B).

**Figure 5.**
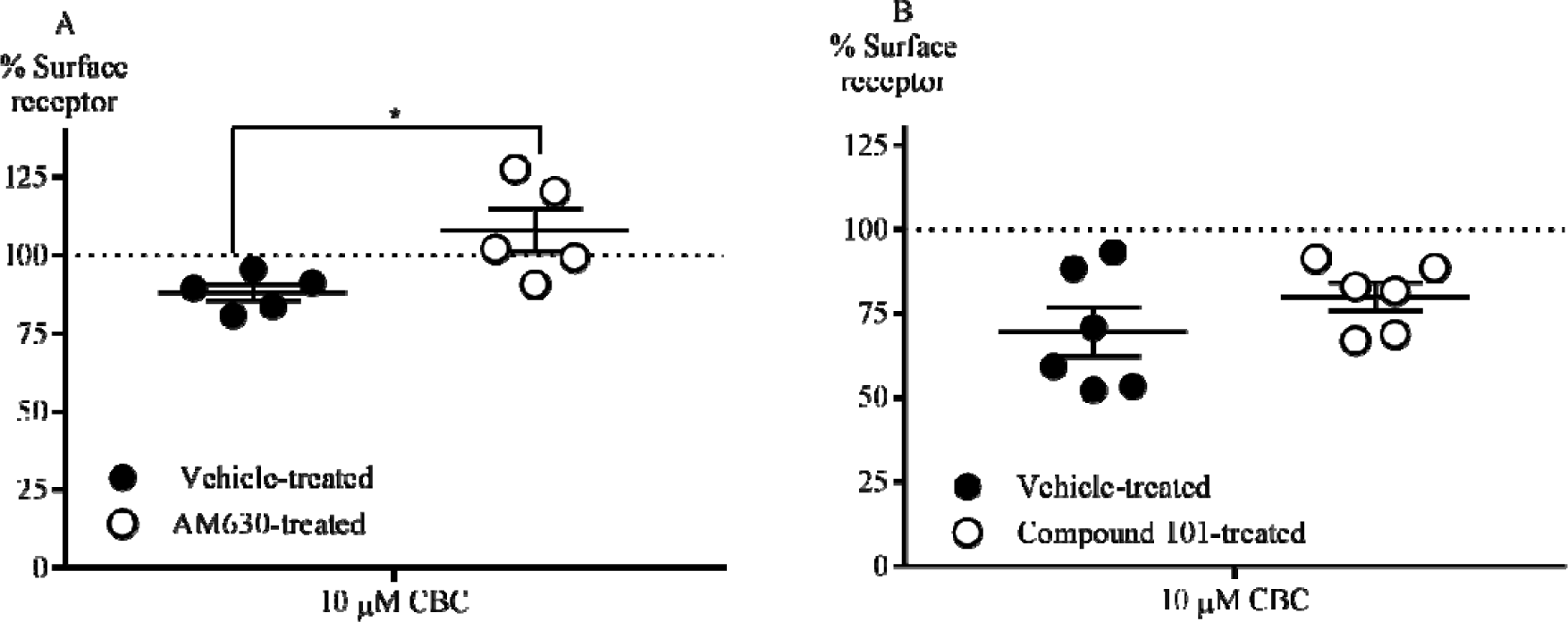
Effect of AM630 and compound 101 on CBC internalisation of CB2 cell surface receptors. (A) Summary data of the effect of AM630 on CBC internationalization of surface receptors. Cells were pre-treated with AM630 (3 µM, 5 mins) followed by CBC (10 µM, 30 min) in the continuous presence of antagonist. (B)Summary data of the effect of compound 101 on CBC internationalization of surface receptors. Cells were pre-treated with compound 101 (10 µM, 60mins), followed by CBC (10 µM, 30 min) in the continuous presence of the GRK2/3 inhibitor. All Results are expressed as mean surface receptor loss ± SEM, which is the percentage difference withvehicle-treated AtT20-CB2 cells, after subtraction of background signal (*n=5-6*).

### Effect of CBC on CB2 signalling desensitisation

In the membrane potential assay, prolonged stimulation of AtT20 cells results in the slow reversal of cellular hyperpolarisation (Cawston et al., 2013). Continuous stimulation of CB2 receptors for 30 minutes by 1 µM CP55,940 or 10 µM CBC resulted in a reversal of the cellular hyperpolarization by 88 ± 3% (Fig 6A) and 73 ± 6%, (Fig 6B), respectively. The desensitisation did not significantly change, when cells were pre-treated with Compound 101 (10 µM, 60mins).

**Figure 6.**
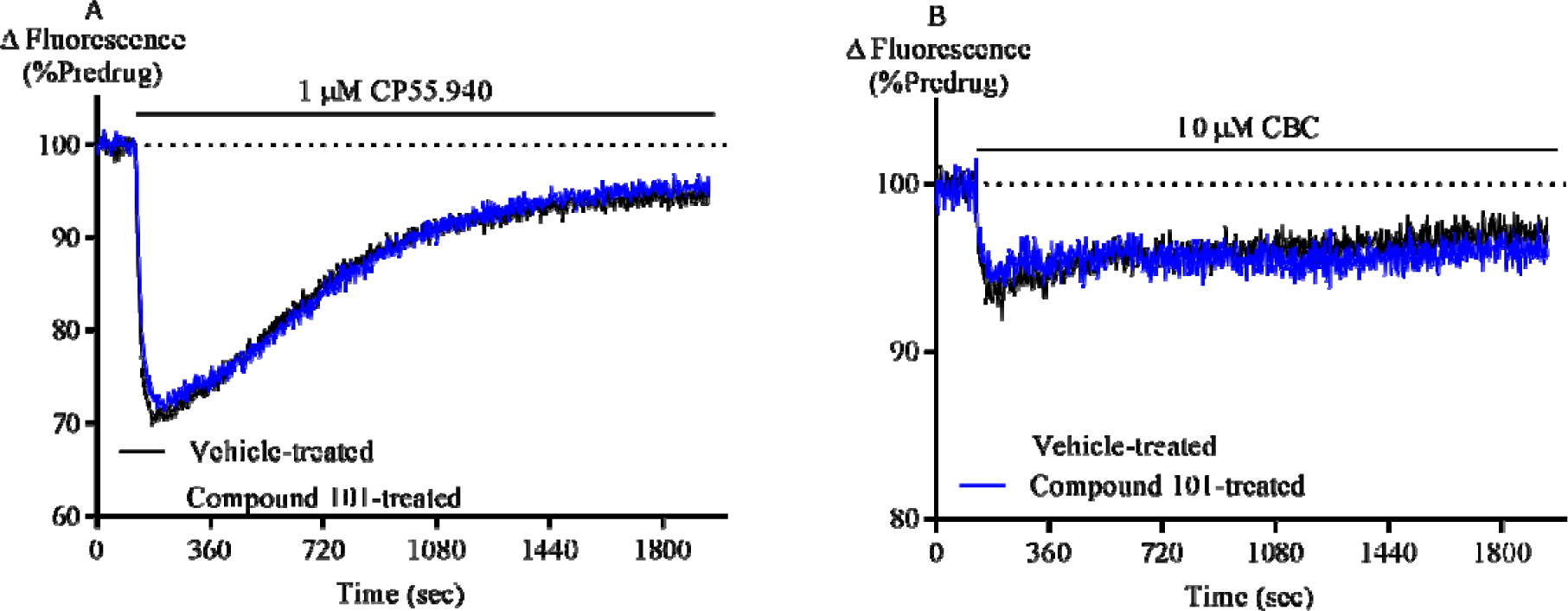
Desensitisation of AtT20-CB2 cells signalling. (A) A representative trace of 1 µM CP55,940 desensitisation of AtT20-CB2 cell signalling in the presence of vehicle or compound 101. (B) A representative trace of 10 µM CBC desensitisation of AtT20 CB2 cell signalling in the presence of vehicle or compound 101. Cell were pre-incubated with compound 101 for 1h (10 µM) before CP55,940 or CBC addition. CP55,940 or CBC were added after 2 mins of baseline reading and read for 30 minutes. (*Note: truncated axes*)

## Discussion

In this study, we have discovered that CBC is a phytocannabinoid with selective CB2 receptor agonist actions. We have also provided evidence that it signals through the Gi/o type G-proteins and induces CB2 receptor internalisation and signalling desensitisation that is independent of GRK2/3 kinases.

CBC produced a dose-dependent cell activation indicated by cellular hyperpolarisation in CB2 cells but with no analogous hyperpolarisation in CB1 cells. This is consistent with a previous finding that CBC apparently does not stimulate [^35^S]GTPγS binding via CB1 expressed in CHO cells (Romano et al., 2013). Although no cannabinoid-antagonist dependent effects have been elucidated in other assays, CBC has been reported to weakly inhibit cellular AEA uptake and the 2-AG hydrolysing enzyme monoacylglycerol lipase (MAGL), both of which may conceivably lead to an indirect activation of cannabinoid receptors through increase in extracellular endocannabinoids (Ligresti, 2006; De Petrocellis et al., 2011). However, the rapid onset of cellular hyperpolarisation in CB2 cells upon addition of CBC suggests a direct receptor activation. Our findings are also consistent with previous studies which concluded that CBC does not significantly contribute to the CB1 receptor mediated psychoactive effects of cannabis *in-vivo* (DeLong et al., 2010; Ilan e al., 2005).

Cannabinoid receptors mediate downstream signalling predominantly through the G_i/o_ protein family (Mallipeddi, et al, 2017), but CB_1_ can couple G_s_-proteins when there is no functional G_i/o_-coupling (Bonhaus et al., 1998; Glass & Felder, 1997), and affect G_q_ in some environments (Lauckner, Hille, & Mackie, 2005). The loss of CBC signalling upon PTX treatment confirms G_i/o_-protein coupling in the hyperpolarization assay, consistent with previous findings with these cells (Banister et al., 2016).

CBC-induced hyperpolarisation in CB2 cells was absent in wild type AtT20 cells and blocked by the selective CB2 receptor antagonist AM630. This blockade is likely due to competitive binding at the CB2 receptor site, supporting the hypothesis that CBC effects are mediated through the CB2 receptor orthosteric site. It is noteworthy that SR144,528, a CB2 antagonist, does not block the anti-inflammatory effects of CBC in both *in vitro* (inhibition of nitrite formation in peritoneal macrophages) and *in vivo* (LPS-induced paw oedema) assays (DeLong et al., 2010; Romano et al., 2013). The receptor mechanisms underlying these anti-inflammatory effects are not yet fully defined.

THC is a low efficacy agonist in many assays of CB1 and CB2 receptor function (Bayewitch et al., 1996; Soethoudt et al., 2017). Therefore, we investigated whether CBC could be acting as an antagonist at the CB1 receptor, since it had been previously reported to bind at the CB1 receptors, albeit with less affinity than CB2 receptors (Rosenthaler et al., 2014). Using sub-maximal concentrations of a high efficacy agonist (300 nM CP55,940), and maximum concentration of a low efficacy agonist (10 μM THC), we found that CBC did not alter the onset, and extent, of cellular hyperpolarisation in CB1 receptor expressing cells. These suggest that CBC does not significantly interact with the CB1 receptor site. In CB2 cells, however, CBC significantly reduced the extent of CP55,940-induced hyperpolarisation.

Stimulation of both CB1 and CB2 receptors have been implicated in antinociception (Bisogno et al., 2009; Guindon, Desroches, & Beaulieu, 2007; Kinsey et al., 2010; La Rana et al., 2006; Lichtman et al., 2004). CB1 receptors are involved in the attenuation of synaptic transmission of nociception in the brain and primary afferent neurons while CB2 contributes to antinociception by inhibiting the release of proinflammatory factors around nociceptive neuron terminals (Manzanares, Julian, & Carrascosa, 2006). Since THC analgesia is at least partly mediated through the CB1 receptors (Mao et al., 2000) and CBC is a ligand for CB2 receptors, it is possible that the potentiation of THC analgesia by CBC, in addition to pharmacokinetic interaction (Davis & Hatoum, 1983; DeLong et al., 2010) may be a result of CBC stimulation of CB2-mediated inhibition of the release of pro-inflammatory factors. Apart from CB2-related anti-inflammatory activities, CBC may act directly or indirectly on proteins such as TRPA1 or adenosine A1 receptors (De Petrocellis et al., 2008; Maione et al., 2011; Shinjyo & Di Marzo, 2013).

Upon sustained exposure to agonists, CB2 receptors undergo receptor internalisation, resulting in signalling desensitisation (Bouaboula, Dussossoy, & Casellas, 1999; Shoemaker et al, 2005). Our results show that CBC caused both loss of surface receptors and signalling desensitisation of CB2 receptors. However, this effect was lesser in magnitude than that observed with CP55,940. This may be due to lower efficacy of CBC in comparison to CP55,940 which is among the most efficacious cannabinoids for internalisation (Atwood et al., 2012). CBC-induced loss of surface CB2 receptors was antagonised by AM630, an effect that further underlines the agonist effect of CBC is CB2 receptor-mediated. AM630 has previously been reported to increase (Grimsey et al., 2011), or have no effect (Atwood et al., 2012), on CB2 surface receptor levels. It is also an inverse agonist at the CB2 receptor (Ross et al., 1999). These imply that AM630-mediated reversal of CBC-induced CB2 internalisation, could be a result of an increase in cell surface receptors by AM630 and not necessarily blockade of CBC internalisation. However, our own observations were that within our experimental conditions, AM630 did not have any appreciable effect on the cell surface receptors. Thus, the effect of AM630 on CBC-induced CB2 internalisation is concluded as a product of direct antagonism at the receptor site.

In the canonical view, signal desensitisation is usually mediated by GPCR kinase (GRK)-mediated phosphorylation of GPCRs; and phosphorylated receptors interact with arrestins to prevent further downstream signalling (Gainetdinov et al., 2004). GPCR kinases involved in CBC-induced CB2 surface internalization and desensitization have not been identified. Our results suggest that GRK2/3 kinases are most likely not involved in these processes. This is in line with previous findings which suggested that GRK2/3 were not likely involved in CB2 internalisation (Bouaboula et al., 1999). Therefore, further investigations are necessary to identify the GPCR kinases involved.

Beta-caryophyllene, which is a terpenoid found in relative abundance within cannabis and food plants, is a naturally occurring CB2-selective agonist (Gertsch et al., 2008). It has both *in vitro* and *in vivo* CB2-mediated anti-inflammatory activities. Here, we have shown that CBC, a phytocannabinoid, is also CB2-selective agonist. This selectivity implies that CBC and/or its derivatives may be further investigated as a potential therapeutic agent that influences the non-psychotropic CB2 receptor pathways of the ECS. Understanding its mechanism of anti-inflammatory activity *in-vitro* and *in-vivo*, as well as activity at other targets, would be valuable in developing its therapeutic potential. Notably, the combination of CBC with THC produces enhanced anti-nociception and anti-inflammatory responses *in vivo* (Davis & Hatoum, 1983; DeLong et al., 2010). This may reflect pharmacokinetic interactions with THC, but also the pharmacodynamic effects of CBC itself on inflammatory processes. Future research might further investigate CBC in combination with THC and cannabidiol to formulate an optimal analgesic cannabis-based medicine, with minor psychotropic effects and potentiated analgesia.

## Acknowledgements

M.U is a recipient of the Macquarie University Postgraduate Research Scholarships for International Students. We thank Lambert Initiative for Cannabinoid Therapeutics for their kind donation of CBC.

## Author contributions

M.U, M.S and M.C designed and analysed experiments. S.D synthesized CBC, M.U conducted all other experiments. M.U, S.D, M.S, I.M and M.C prepared the manuscript. All authors have seen the final paper.

## Abbreviations

2-AG: 2-arachidonoyl-glycerol;
AEA: anandamide;
AtT20-CB1: Mouse pituitary tumour cells stably transfected with HA human CB1 cells;
AtT20-CB2: Mouse pituitary tumour cells stably transfected with HA human CB2 cells;
CB1: Cannabinoid receptor type 1
CB2: Cannabinoid receptor type 2;
CBC: cannabichromene;
CBD: Cannabidiol;
CBN: Cannabinol;
ECS: Endocannabinoid system;
GRK: G-protein coupled receptor kinase;
HA: Haemagglutinin
PTX: Pertussis toxin;
THC: Tetrahydrocannabinol;
THCV: Tetrahydrocannabivarin.

